# High-Throughput Single-Cell Spectroscopy Using Phasor Analysis of Spectral Flow Cytometry

**DOI:** 10.64898/2026.02.27.708623

**Authors:** Bruno Pannunzio, Paula Céspedes, Marcela Díaz, Daniela Alí, Analía Rial, Leonel Malacrida

**Affiliations:** Advanced Bioimaging Unit, Institut Pasteur de Montevideo-Universidad de la República, Uruguay; Unidad Académica de Histología y Embriología, Facultad de Medicina-Universidad de la República-Uruguay.; Laboratorio de Genética Molecular Humana, Institut Pasteur de Montevideo, Uruguay; Unidad Académica de Desarrollo Biotecnológico, Facultad de Medicina-Universidad de la República-Uruguay.; Unidad Académica de Fisiopatología, Hospital del Clínicas, Facultad de Medicina-Universidad de la República-Uruguay.

## Abstract

Phasor analysis is a well-established tool in hyperspectral and lifetime microscopy, providing a powerful, fit-free approach for interpreting complex fluorescence. However, its application has remained largely restricted to imaging-based modalities. Spectral flow cytometry (SFC) enables acquisition of full emission spectra from large numbers of independent single-cell events, offering superior statistical power compared to microscopy, albeit at the expense of spatial and temporal information. Here, we present the first implementation of spectral phasor analysis for SFC (phSFC), establishing a unified analytical framework that preserves interpretative continuity with hyperspectral microscopy while extending phasor-based analysis to high-throughput, single-cell measurements. Using the membrane-sensitive probe LAURDAN as a benchmark, we demonstrate that SFC reproduces phasor signatures of membrane order previously reported by hyperspectral confocal microscopy (HSI). We performed comparative analyses using multilamellar lipid vesicles (MLVs) prepared from known physical order compositions. Both modalities, SFC and HSI, accurately resolved MLVs with fluid, gel and liquid-ordered and liquid-disorder membrane phases, capturing cholesterol-dependent spectral shifts, including trajectories associated with mixtures of the different lipid phase behavior. Although absolute phasor coordinates differed between modalities due to distinct spectral sampling and detector configurations, the relative organization of membrane physical states was preserved. Notably, SFC produced more compact phasor distributions, consistent with larger sample size and enhanced statistical robustness. To further extend phSFC, we first evaluated its capacity to resolve membrane changes in live cultured cells following cholesterol depletion, establishing consistency between HSI and SFC measurements. We then applied phSFC to detect membrane dynamics in primary leukocytes isolated from bronchoalveolar lavage of mice with inflammation-associated lung pathology. LAURDAN fluorescence in the presence of autofluorescence and antibody-derived signals is quantified and discussed with simple solution by n-harmonic phasor analysis unmixing. Together, these results establish SFC as a robust and complementary extension of LAURDAN phasor analysis, bridging HSI and high-throughput flow cytometry measurements.

## Introduction

Phasor analysis has emerged as a powerful framework for the fit-free analysis of spectral fluorescence data, enabling intuitive and quantitative decomposition of complex emission signals in the Fourier domain (Fereidouni et al., 2012). By mapping spectral information into a compact graphical space, phasor analysis circumvents the need for explicit spectral models or prior assumptions and has become widely adopted in fluorescence microscopy for applications ranging from fluorescence lifetime imaging (FLIM) to hyperspectral analysis (HSI) (Malacrida, 2023; Malacrida et al., 2021). Despite its success in imaging-based modalities, this approach has not yet been translated to spectral flow cytometry (SFC), a high-throughput technique that captures the full fluorescence emission spectra across multiple detectors for large populations of independently sampled particles at rates unattainable by microscopy (Nolan, 2022; Paul Robinson and Rajwa, 2024). Extending phasor analysis to SFC would provide a fundamentally new way to interrogate cellular spectral signatures at scale, but requires careful adaptation to flow-based acquisition and data structures.

SFC offers complementary valuable information to microscopy for spectral analysis. Its high-throughput nature enables statistically robust characterization of heterogeneous cell populations, facilitating the detection of rare subsets and subtle spectral shifts that are impractical to resolve by imaging alone. Importantly, the multidimensional data architecture of flow cytometry, which integrates spectral fluorescence information with light-scatter and conventional fluorescence parameters, aligns naturally with the phasor approach and enables seamless integration of spectral analysis with established immunophenotyping or reporter-based assays.

Current analysis pipelines in SFC typically rely on spectral unmixing based on reference controls and manual gating strategies defined by intensity thresholds, which are highly user-dependent (Aghaeepour et al., 2013; Andronico et al., 2025; Nolan, 2022). While effective in many contexts, these methods have inherent limitations. Spectral unmixing requires accurate reference spectra, averaging out poor components (below 5%), and is unable to consider autofluorescence as a component part of the fluorescence recorded, while threshold-based gating lacks the precision required to accurately capture continuous or overlapping spectral changes (Mage et al., 2025; Robinson, 2022; Schmutz et al., 2016). In contrast, phasor analysis provides a reference-free and unbiased representation of spectral data, preserving its continuous nature and enabling direct visualization of spectral signatures without compensation (Fereidouni et al., 2012). This reduces user-dependent bias and facilitates the detection of gradual or mixed spectral states. The model-free nature of phasor plots makes this approach ideal for solvatochromic probes and endogenous fluorescence.

By retaining spectral continuity, phasor analysis enables the identification of trajectories and intermediate states that are often obscured by conventional discrete classification approaches. Such capabilities are particularly relevant for biological systems characterized by gradual transitions, including metabolic remodeling, membrane reorganization, or responses to pharmacological or inflammatory stimuli (Malacrida et al., 2021). Moreover, phasor representations offer intrinsic quality control, since spectra affected by noise or experimental artifacts naturally segregate as outliers in phasor space, enabling rapid identification without additional preprocessing. These features position phasor analysis as a complementary and, in some cases, superior analytical paradigm for high-dimensional SFC data.

In this work, we present the first implementation of phasor-based SFC (phSFC), establishing a general framework for fit-free, high-throughput spectral characterization at the single-cell level. An overview of the analytical framework used to adapt spectral phasor analysis to SFC, and to relate it to HSI-based approaches. As a benchmark application, we focus on LAURDAN fluorescence, an environmentally sensitive probe used for studies of membrane dynamics. LAURDAN changes the emission spectrum in response to the physical state of the membranes (Díaz et al., 2026; Malacrida et al., 2016, 2015, 2014). Using multilamellar lipid vesicles with defined physical states and increasing cholesterol fraction, we demonstrate that phSFC data faithfully reproduces established trends previously observed by HSI and phasor analysis. Direct comparison of phasor representations derived from HSI and SFC confirms strong agreement in spectral organization, while highlighting differences arising from acquisition modality and sampling statistics.

We further extend this approach to living systems by analyzing LAURDAN-stained cells under conditions of controlled membrane perturbation. Using methyl-β-cyclodextrin (MβCD)–treated cells to reduce membrane cholesterol, we show that SFC captures membrane-order–dependent phasor shifts consistent with prior microscopy-based studies (Díaz et al., 2026), while providing enhanced population-level resolution. Finally, to demonstrate applicability in a physiologically relevant *in vivo* context, we apply phSFC approach to leukocytes isolated from bronchoalveolar lavage following lipopolysaccharide (LPS) challenge in mice.

Together, these results demonstrate that phasor-based spectral flow cytometry (phSFC) enables robust, reference-free characterization of membrane organization across model systems, cultured cells, and *in vivo* derived primary cells, bridging the gap between microscopy-based spectral analysis and high-throughput cytometric measurements.

## Materials and methods

### LAURDAN stain preparation

The membrane-sensitive dye LAURDAN (6-dodecanoyl-2-dimethylaminonaphthalene; Sigma-Aldrich #40227) was dissolved in dimethyl sulfoxide (DMSO) to prepare a 3.1 mM stock solution. The concentration was determined spectrophotometrically using a Jasco V-750 spectrophotometer with a molar extinction coefficient of 20,000 ML¹ cmL¹ (at 364 nm in methanol).

### Preparation and Staining of Multilamellar Vesicles (MLVs) with LAURDAN

Multilamellar vesicles (MLVs) exhibiting fluid, solid, or liquid-order/disorder phases were prepared using the following lipids: 1,2-dioleoyl-sn-glycero-3-phosphocholine (DOPC; Avanti Polar Lipids, LLC #850375C), 1,2-dipalmitoyl-sn-glycero-3-phosphocholine (DPPC; Avanti Polar Lipids, LLC #850355), and an increasing concentration of cholesterol (Avanti Polar Lipids, LLC #700000). Stock solutions of each lipid were prepared in chloroform and mixed to obtain three lipid compositions at a total phospholipid concentration of 0.5 mM: (i) DOPC, (ii) DPPC, and (iii) a 1:1 molar mixture of DOPC: DPPC. For each lipid composition, cholesterol was added at three molar ratios (Phospholipid: Cholesterol) of 0, 2:1 (33% cholesterol), and 1:1 (50% cholesterol). The lipid mixtures were dried under a gentle stream of N_2_ gas and subsequently placed under vacuum overnight to remove solvent residuals. The resulting lipid films were rehydrated in a preheated 10 mM HEPES buffer pH 7.4, followed by three cycles of vortexing and heating at 45 °C to promote MLVs formation. The LAURDAN was added to the lipid mixture before hydration at a final dye-to-lipid molar ratio of 1:200. For imaging, aliquots of each suspension were deposited onto glass-bottom Petri dishes and allowed to settle for 1 h before acquisition. All images were acquired at 21 °C.

### Vero Cell Culture and Methyl-**β**-Cyclodextrin (M**β**CD) Treatment

The protocol was performed as previously described by (Díaz et al., 2026). Briefly, Vero cells were cultured in Dulbecco’s Modified Eagle Medium (DMEM) supplemented with 10% Fetal Bovine Serum (FBS) at 37°C under 5% COL. For cell staining, a 15 µM final concentration of LAURDAN in DMEM was used with the cells and incubated for 60 min at 37°C before imaging. For cytometry, the monolayer was detached with 0.25% trypsin, diluted 1:10 in DMEM, and transferred to a 35-mm glass-bottom Petri dish for overnight adherence. Membrane cholesterol levels were modulated by treating cells with methyl-β-cyclodextrin (MβCD; Sigma-Aldrich, #C4555). The MβCD was prepared in serum-free DMEMA at 38 mM stock solution. Then cells were incubated with 10 mM MβCD for 90 minutes at 37 °C. After treatment, cells were washed three times with PBS and subsequently labeled with 15 µM final concentration of LAURDAN in serum-free DMEM for 30 minutes before imaging. For SFC analyses, cells were detached using trypsin and resuspended in serum-free DMEM containing 15 µM final concentration of LAURDAN.

### Animal model and leukocytes obtained from Bronchoalveolar Lavage (BAL)

CD1 male mice (6-8 weeks old) were anesthetized by intraperitoneal injection 2.2 mg ketamine plus 0.11 mg xylazine in a total volume of 200 µl and then intranasally instilled with LPS (Escherichia coli; O55:B5 serotype; Sigma Aldrich) at a dose of 4 mg/kg body weight (45 ul/dose) or 45 ul of saline (Burkard et al., 2023). After 18 hours, mice were sacrificed by cervical dislocation, and bronchoalveolar lavages (BAL) were performed as previously described (Suárez et al., 2020). The BAL fluid was centrifuged (400g, 10min, 4°C), washed 2 times with PBS-containing 1mM Ethylenediaminetetraacetic acid (EDTA) and 2% fetal bovine serum (Capricorn, FACS-DTA) and finally resuspended in FACS (Fluorescence-Activated Cell Sorting) buffer with EDTA. For immunophenotyping of bronchoalveolar lavage (BAL) cells, 1 × 10L cells per tube were seeded and stained with eFluor 660–conjugated anti-mouse CD11c (clone N418, eBioscience) and APC-Cy7–conjugated anti-mouse CD45 (clone 30-F11, BioLegend), in the presence of 15 µM of LAURDAN. All animal experimentation protocols were approved by the University’s Ethical Committee for Animal Experimentation, Uruguay (protocol number: 2425).

### Spectral Flow Cytometry (SFC)

Flow cytometry data were acquired using a Cytek Aurora spectral flow cytometer (Cytek Biosciences) equipped with three lasers (405 nm, 488 nm, and 633 nm). Before each acquisition session, the instrument was allowed to warm up for 30 min, and daily quality control was performed to ensure laser stability and optimal spectral performance using SpectroFlo QC calibration beads (1 drop in 0.3–0.5 mL sheath fluid; lot 2005). Data acquisition was performed at a low flow rate for MLVs and at a medium flow rate for VERO cells, acquiring 20,000 and 10,000 events per sample, respectively. Given that MLV signals are detected close to the electronic noise threshold of the instrument, additional controls were performed to validate the correct identification of the MLV population. These included acquisition of HEPES buffer alone, a 4X Triton solution alone, and MLVs treated with 4X Triton. These controls allowed discrimination between electronic noise, detergent-induced lipid solubilization, and potential aggregates, confirming that the gated events corresponded to lipid vesicles rather than noise or debris. These controls are shown in Supplementary Figure 1.

### Hyperspectral Imaging (HSI) Acquisition

HSI was performed using a Zeiss LSM 880 confocal microscope (Zeiss, Oberkochen, Germany) equipped with a 63× oil-immersion objective (NA 1.4, Plan-Apochromat, DIC M27, Zeiss). LAURDAN was excited using a 405 nm laser diode, and emission spectra were collected in lambda mode using Zen Black 2.3 software (Zeiss). A gallium arsenide phosphide photomultiplier tube (GaAsP-PMT, 650V master gain) was used to sequentially record fluorescence across 28 spectral channels (between 423-693 nm, with 10 nm steps). Images were acquired at a resolution of 512 × 512 pixels, with a pixel dwell time of 0.77 µs and a pixel size of 105 nm.

### Data Analysis Framework

All data processing and analysis were performed using custom-written Python code developed specifically for this study, following a unified analytical framework adapted to the distinct data structures of spectral flow cytometry and hyperspectral confocal microscopy (Figure 1). The workflow was designed to ensure conceptual continuity between modalities while allowing modality-specific preprocessing steps. All analyses were executed in Python (version 3.13.9) in a reproducible computational environment.

**Figure 1.**
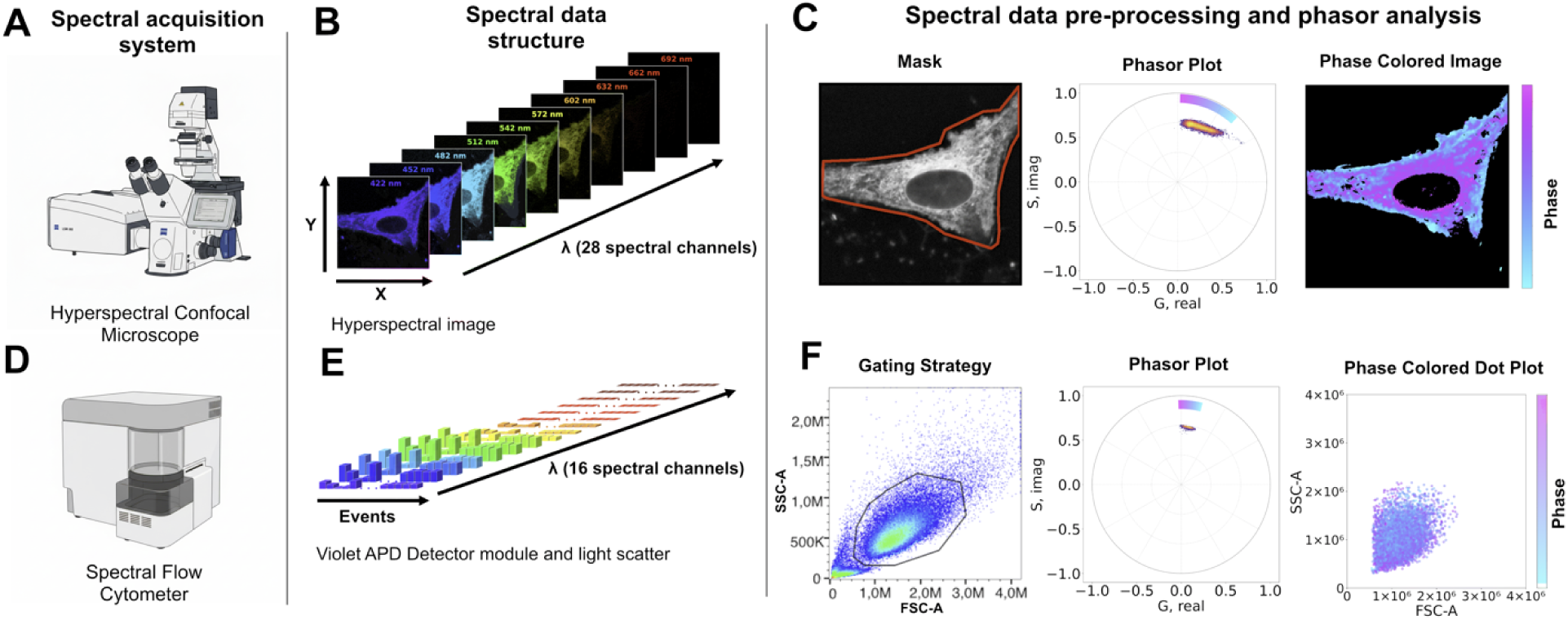
Overview of the spectral phasor analytical framework for SFC and HSI. The top row **(A–C)** shows the workflow for SFC, and the bottom row **(D–F)** the corresponding workflow for HSI. **(A, D)** Spectral acquisition platforms: Cytek Aurora flow cytometer **(A)** and Zeiss LSM 880 confocal microscope **(D)**. **(B, E)** Data structure generated by each modality: flow cytometry produces event-based spectra across 16 violet channels (λ) per independently sampled event (2-D stack), whereas microscopy generates spatially resolved spectral 3D-stacks (X, Y, λ). **(C, F)** Modality-specific preprocessing followed by Fourier-based computation of phasor coordinates (g, s). In flow cytometry, gated events (single cells) are transformed, and phasor-derived parameters can be mapped onto conventional cytometry plots; in microscopy, every pixel (after masking) are transformed and phasor-derived parameters can be mapped back into the image to visualize spatial spectral variations.

### Spectral Flow Cytometry Data Processing

Raw SFC data were acquired as “.fcs” files and initially processed using FlowJo software (BD Biosciences) for population selection. Because SFC measurements consist of independently sampled fluorescence spectra accompanied by light-scatter parameters, population selection was performed prior to spectral phasor analysis to isolate biologically relevant measurements and exclude debris or aggregates. Gating strategies were tailored to each sample type while following a common hierarchical approach. For MLVs and VERO cells, populations were first defined using side-scatter area (SSC-A) versus forward-scatter area (FSC-A), followed by FSC-A versus FSC-H to exclude aggregates and select single particles or cells. LAURDAN-positive events were identified based on fluorescence intensity in the V3 channel. For BAL samples, an additional immunophenotyping step was incorporated before LAURDAN-based selection. After initial gating on FSC-A versus SSC-A and FSC-A versus FSC-H to exclude debris and doublets, CD45-positive events were identified using APC-Cy7 fluorescence (R7 channel). CD45L cells were subsequently gated for CD11c expression using eFluor-660 fluorescence (R2 channel).

Following gating, the selected populations were exported from FlowJo as separate “.fcs” files and used as input for spectral phasor analysis. Exported files were read into Python using the fcsparser library (Yurtsev, 2026), which was used to extract spectral fluorescence intensities, light-scatter parameters, and associated metadata. For spectral phasor analysis, only the 16 fluorescence channels excited by the 405 nm (violet) laser (V1 to V16 channels, 428 to 812 nm wavelength range), corresponding to the LAURDAN emission spectrum, were selected. Spectral data were assembled into two-dimensional arrays defined by independently sampled measurements and wavelength channels (Figure 1B, top row).

### Hyperspectral Microscopy Data Processing

Hyperspectral confocal microscopy data were acquired as “.lsm” files and imported into Python using PhasorPy v0.9 (Gohlke et al., 2026), which provides native support for multi-dimensional spectral image data. Spectral image stacks were organized as three-dimensional arrays defined by spatial coordinates (X, Y) and wavelength channels (Figure 1B, bottom row). To restrict analysis to relevant regions, an interactive region-of-interest selection procedure was implemented using Matplotlib (Hunter, 2007). Average intensity images were displayed, and binary masks were manually defined to isolate lipid vesicles or cellular regions while excluding background. These masks were applied to the corresponding spectral image stacks, ensuring that only the selected pixels contributed to subsequent phasor analysis. Additional preprocessing steps were applied using tools available in the PhasorPy library (https://www.phasorpy.org), including intensity thresholding for background removal and noise filtering using three iterations of a 3×3 median filter.

### Spectral Phasor and Component Analysis

Spectral phasor transformation and component analysis were performed using PhasorPy for both SFC and HSI datasets. To map the spectral data into the phasor domain, each spectral vector (representing either an individual measurement in flow cytometry or a single pixel in hyperspectral microscopy) was processed via a Fourier-based transformation. This method calculates the real (G) and imaginary (S) components of the *n^th^* harmonic to define the corresponding phasor coordinates as follows (Fereidouni et al., 2012):

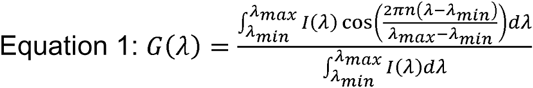

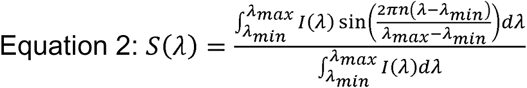

Where *I(*λ) represent the spectral intensity measured at a specific wavelength λ within the detection boundaries defined by λ*_min_* and λ*_max_*. The variable n denotes the harmonic number. The denominator ∫ *I(λ)dλ*. acts as a normalization factor to ensure that the calculated phasor coordinates remain independent of the total signal intensity.

Reference phasor coordinates for liquid ordered and disordered LAURDAN emission were defined using measurements from lipid vesicles with knowing membrane physical states, separately for each acquisition modality to account for differences in spectral range and detector configuration. For experiments involving additional fluorescent contributors, including autofluorescence and antibody-derived signals, reference phasor coordinates were defined from control measurements. Multi-component phasor analysis using 2 harmonics were then applied to estimate fractional contributions of each component on a per-measurement or per-pixel basis.

### Visualization and Statistical Analysis

Phasor plots, histograms, and back-projected visualizations were generated using Matplotlib and the PhasorPy library. For flow cytometry data, phasor-derived parameters were mapped back onto conventional dot plots, enabling integrated visualization of spectral properties alongside scatter and marker-based parameters. For microscopy data, phasor parameters were back-projected into spatial images to generate false-color representations of local spectral properties.

Statistical summaries and data organization were performed using Pandas library (McKinney, 2010). Distribution-based visualizations, including violin plots, were generated using the SuperViolin plotting library (Kenny and Schoen, 2021). All figures were generated directly from the analysis scripts to ensure consistency between quantitative results and visual representations.

## Results

### Consistency Between SFC and HSI in LAURDAN-Based Membrane Fluidity Assessment

The primary objective of this study was to assess whether SFC can reproduce the well-established LAURDAN-based membrane fluidity measurements obtained by HSI. To this end, we compared the phasor distributions of LAURDAN-stained multilamellar vesicles (MLVs) acquired using both techniques.

Hyperspectral microscopy analysis of DOPC (fluid phase) and DPPC (solid phase) MLVs yielded the expected separation in phasor space, with DOPC MLVs clustering in the high-fluidity region and DPPC MLVs occupying the ordered membrane region (Figure 2). These results were consistent with numerous previous studies (Bacalum et al., 2024; Kaiser et al., 2009; Ma et al., 2018; Malacrida et al., 2016, 2014) that have established microscopy-based LAURDAN phasor analysis as a reliable method for membrane order assessment. Notably, when we analyzed the same MLV preparations using SFC, we observed highly similar clustering patterns in phasor space (Figure 2). The SFC data yielded phasor distributions for both DOPC and DPPC MLVs that fell within the regions expected from microscopy-based membrane fluidity measurements. Differences in the exact cluster positions were observed, consistent with the distinct spectral ranges and channel configurations of the two acquisition systems.

**Figure 2.**
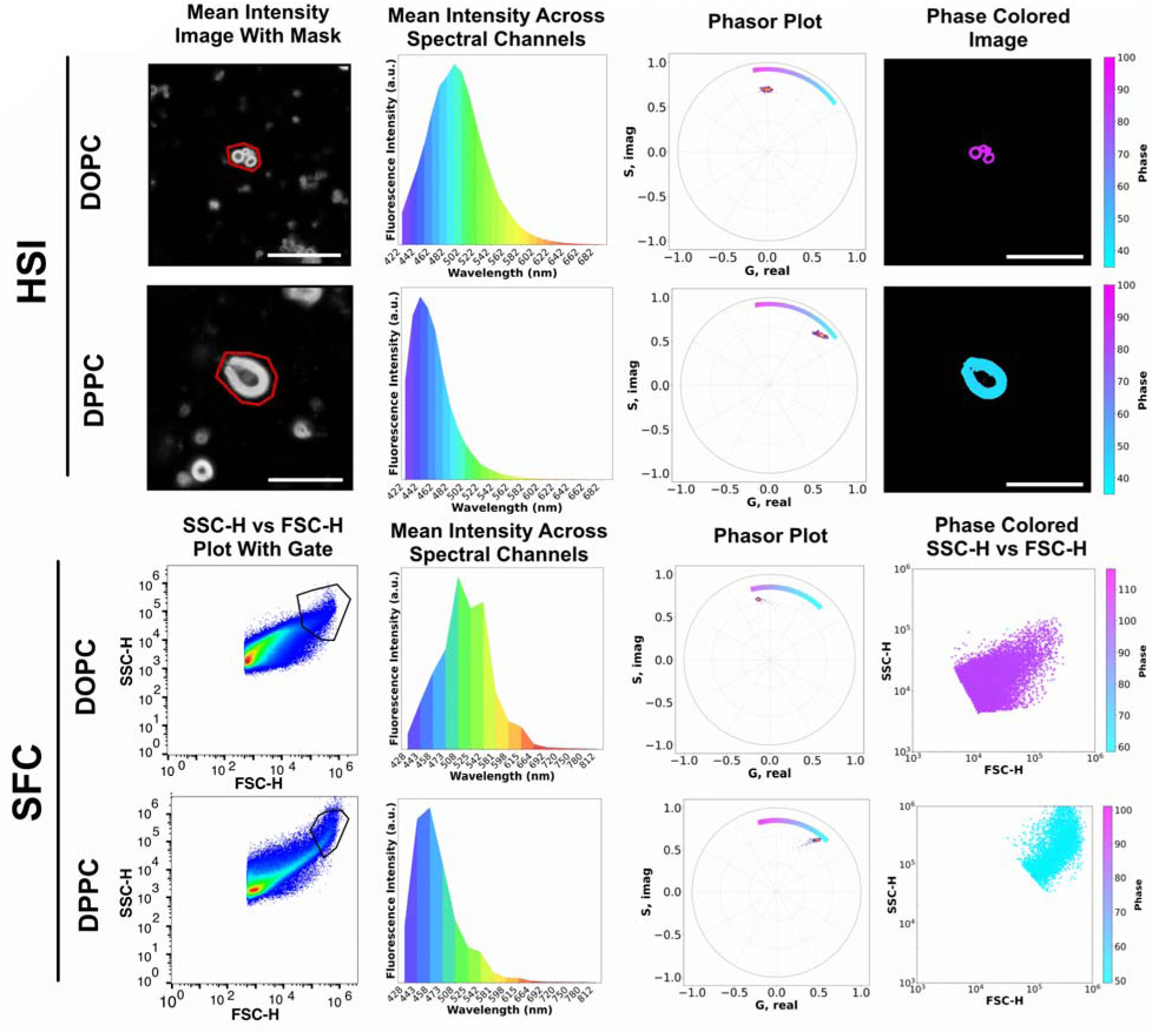
Spectral Phasor Analysis Workflow for Multilamellar Vesicles (MLVs) using HSI and SFC. Comparative analysis workflow for assessing membrane order in DOPC or DPPC multilamellar vesicles stained with LAURDAN, using HSI (top rows) or SFC (bottom rows). The microscopy workflow starts with masking (red outline) using the average intensity image of the MLVs, followed by the average emission signal across 28 spectral channels. Then, data is transformed to generate a phasor plot. We use a phase range (spectral center of mass) color bar to highlight spectral shift. The final step is the average intensity image color mapped to show the phase angle distribution across the imaged MLVs. The scale bars in the intensity and phase-colored images correspond to 20 µm. In the SFC workflow, the analysis begins with an FSC-A versus SSC-A dot plot to gate the MLV population, followed by the average emission signal across all 16 spectral channels. Then, data transformation into the phasor plot was calculated, including the associated phase range color bar (blue is the smallest phase and red the largest phase values). Finally, the original flow cytometry events (dot plot) are colored according to their calculated phase angle.

To further assess the agreement between microscopy- and flow cytometry–based spectral phasor measurements across a broader range of membranes with different lipid mixtures, phasor distributions were compared for DOPC, DPPC, and DOPC/DPPC MLVs prepared with increasing cholesterol concentrations (Figure 3). Across both acquisition modalities, the relative organization of the different lipid mixtures in phasor space were preserved. In both microscopy-and cytometry-derived phasor plots, LAURDAN in DOPC/DPPC with 0% cholesterol MLVs fall in the line that join the pure components, gel (DPPC, blue LAURDAN emission) and fluid (DOPC, green LAURDAN emission) membrane. This result is indicative of the additive properties of phasor, and its capability to quantify lipid fraction by simple linear algebra at the phasor space. Interestedly, increasing concentration of cholesterol (33 and 50 %) in DOPC MLVs exhibit LAURDAN spectral toward blue emission in a trajectory, indicative of a phase transition from fluid to liquid-disorder state.

**Figure 3.**
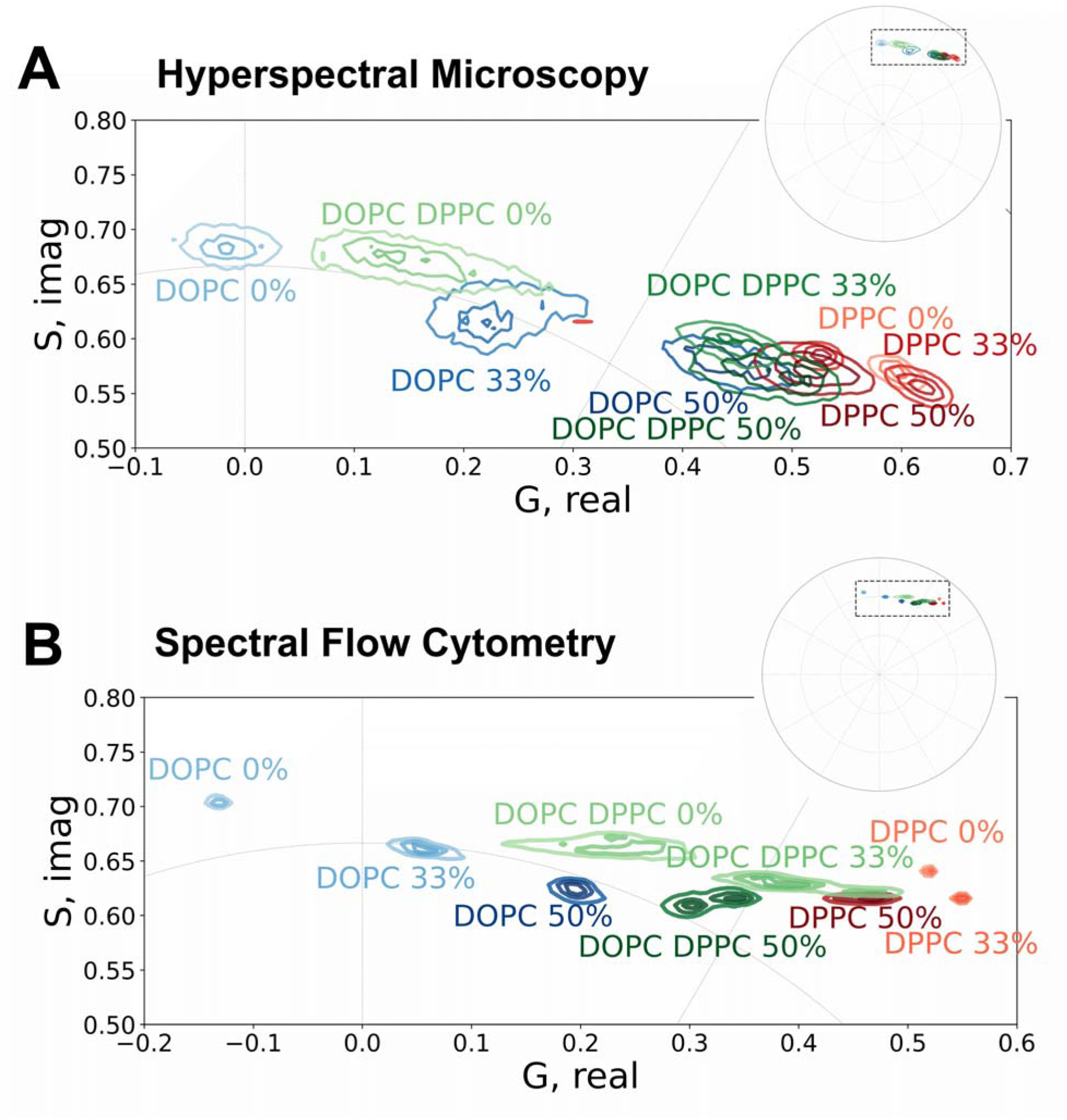
Comparative spectral phasor analysis of multilamellar vesicles (MLVs) by SFC and HSI. Spectral phasor plots for six MLV conditions acquired by HSI (**A,**) and SFC (**B**). The conditions include DOPC, an equimolar DOPC/DPPC mixture, and DPPC, each prepared with three cholesterol concentrations (0%, 33%, and 50%). Data are shown as contour plots in phasor space, with DOPC conditions in blue, DOPC/DPPC mixtures in green, and DPPC conditions in red. For clarity, only the first three contour levels are displayed to highlight overlapping clusters. The mean center of each distribution is indicated by a dot in the corresponding color. Phasor distributions represent merged data from *n* = 2 independent MLV preparations per condition. Each panel displays a magnified view of the phasor region indicated by the dashed rectangle in the full phasor plot inset (top right).

In contrast, DPPC and DOPC/DPPC MLVs did not display a straight linear combination upon increasing concentration of cholesterol. Cluster positions for increasing cholesterol concentrations show a more convoluted transition for membranes moving from pure gel or mixture fluid/gel. For these conditions, changes in cholesterol content resulted in spectral shifts in phasor space that deviated from a simple linear combination of two states. Notably, cholesterol-enriched MLV membranes are located closer to the center of the phasor plot. Cholesterol shifts the LAURDAN phasor coordinates toward the center (reduced modulation), indicating spectral broadening consistent with a more heterogeneous environment for LAURDAN.

At each cholesterol concentration, the phasor distributions of the DOPC/DPPC mixtures were consistently positioned between those of the corresponding pure DOPC and pure DPPC conditions containing the same cholesterol level, in both HSI and SFC measurements. This fact is indicative of membrane coexistence at different fraction of liquid-order and liquid-disorder at MLVs membranes.

Phasor distributions obtained by SFC were more compact than those derived from HSI. This result was expected and reflects the distinct nature of the information provided by each modality. In HSI, each point in phasor space corresponds to an individual pixel from the original image, thereby preserving spatial heterogeneity within single cells. In contrast, in SFC each point represents an independently sampled event, typically a single MLV or cell, whose fluorescence signal is integrated across the entire event. Consequently, spectral variability that appears as a dispersed distribution for a single MLV/cell in HSI is averaged into a single phasor coordinate in SFC. This intrinsic signal integration results in reduced dispersion and tighter clustering in phasor space for SFC, together with differences in signal-to-noise characteristics between the two approaches.

### Comparison of Linear Combination Relationships Across Lipid and Cholesterol fraction by HSI and SFC

To further compare how linear combination relationships are captured by HSI and SFC, the same datasets were re-represented to explicitly evaluate the agreement between measured phasor distributions and the values predicted from component fraction analysis (Figure 4). For DOPC MLVs, the phasor distributions corresponding to 0% and 50% cholesterol were used to define the endpoints of the linear combination, and the distribution at 33% cholesterol was evaluated relative to the expected intermediate position.

**Figure 4.**
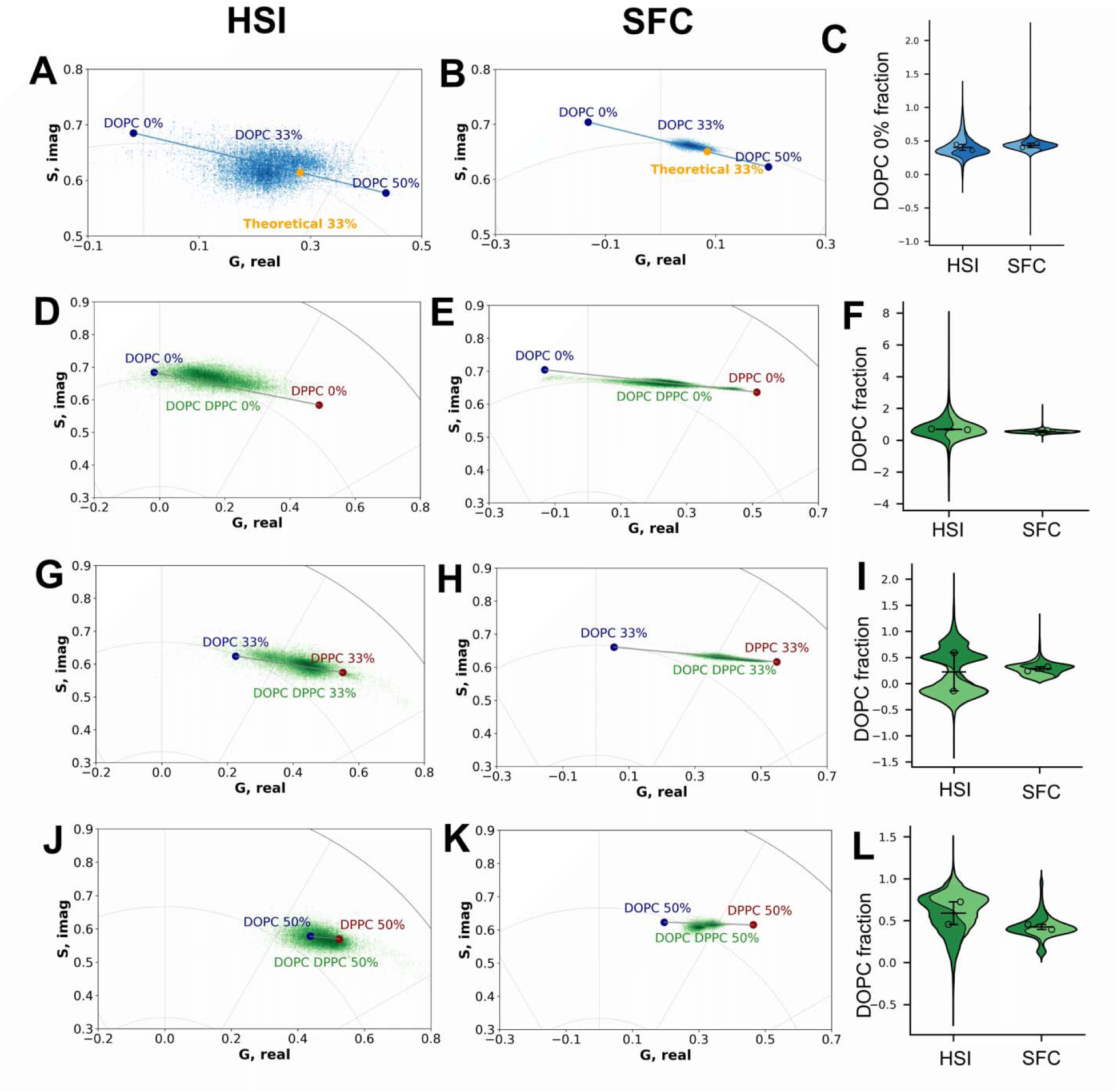
Linear combination analysis of spectral phasor data across lipid mixtures and cholesterol fraction in MLVs measured by HSI and SFC. Spectral phasor–based linear combination analysis derived from the MLV datasets shown in Figure 3. Panels are organized by cholesterol content within the equimolar DOPC/DPPC mixture, with HSI, SFC, and fraction-based quantification shown sequentially. (**A**) HSI phasor representation for pure DOPC MLVs at 0%, 33%, and 50% cholesterol. Mean phasor centers of the 0% and 50% cholesterol conditions are shown as dots, while the 33% condition is displayed as a two-dimensional density plot, illustrating its intermediate position along the linear combination trajectory. (**B**) Corresponding SFC phasor representation for pure DOPC MLVs. (**C**) Violin plot showing the calculated fractional contribution of the 0% cholesterol DOPC component. Each violin represents pooled pixel- or event-level fractions, with individual segments corresponding to independent experiments (*n* = 2). **(D–F)** Calculation of the DOPC fractional contribution within equimolar DOPC/DPPC MLVs at 0% cholesterol. **(D)** HSI phasor representation, showing the DOPC/DPPC mixture positioned relative to the pure DOPC and DPPC reference states. **(E)** Corresponding SFC phasor representation. **(F)** Violin plot of the estimated DOPC fraction for the 0% cholesterol condition, displayed and quantified as described in panel C. **(G–I)** Same analysis as in panels D–F for equimolar DOPC/DPPC MLVs at 33% cholesterol. **(J–L)** Same analysis as in panels D–F for equimolar DOPC/DPPC MLVs at 50% cholesterol.

In the case of SFC, the phasor distribution for DOPC at 33% cholesterol (Figure 4E) was closely centered around the theoretical intermediate value predicted from the linear combination of the 0% and 50% cholesterol conditions, with the mean and spread of the distribution largely confined within the expected range. In contrast, while HSI measurements also showed an intermediate distribution for 33% cholesterol (Figure 4A), the corresponding phasor values exhibited a broader deviation from the predicted mean position.

A similar trend was observed for mixed DOPC/DPPC MLVs across cholesterol concentrations. For each cholesterol level (0%, 33%, and 50%), the phasor distributions of the mixtures measured by SFC (Figure 4F–H) were closely aligned with the expected linear combination between the mean centers of the corresponding pure DOPC and DPPC conditions than those obtained by HSI (Figure 4B–D). Across all mixture conditions, cytometry-derived phasor distributions were centered nearer to the predicted values and displayed reduced dispersion relative to microscopy-derived distributions.

### Evaluation of Live-Cell Membrane Properties Using LAURDAN Phasor Analysis by HSI and SFC

To extend LAURDAN spectral phasor analysis from model membranes to a biologically relevant system, live Vero cells stained with LAURDAN were analyzed by HSI and SFC under control conditions and following methyl-β-cyclodextrin (MβCD) treatment (Figure 5). Cholesterol-dependent shifts in LAURDAN fluorescence induced by MβCD have been previously demonstrated for this cellular model using HSI. The incubation with MβCD led to a reproducible green displacement of LAURDAN fluorescence at the phasor distribution associated with changes in membrane order (Díaz et al., 2026). Building on this prior work, the present study evaluates whether comparable shifts can be detected using SFC.

**Figure 5.**
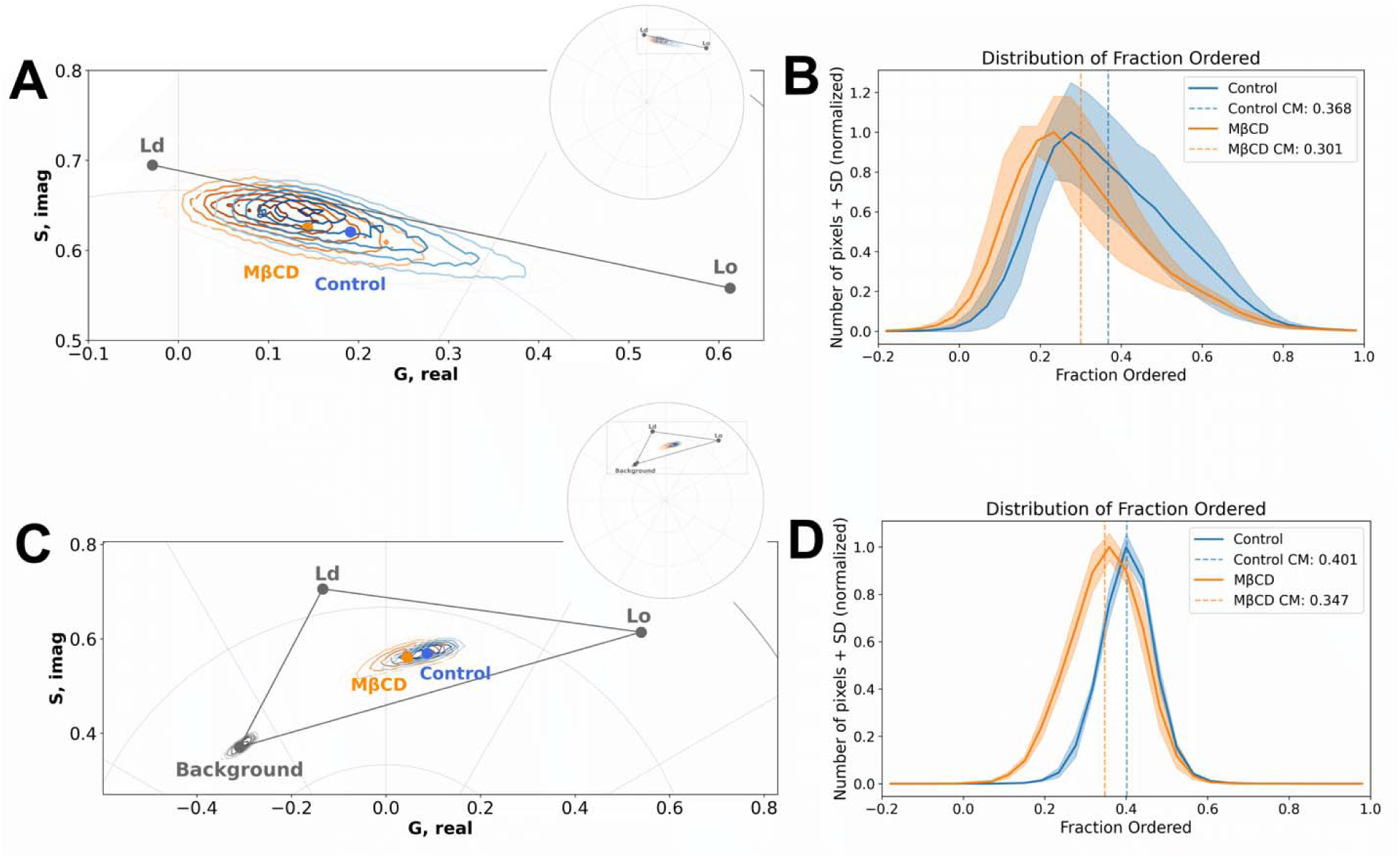
LAURDAN spectral phasor analysis of membrane order in live Vero cells by HSI and SFC. **(A–B)** HSI measurements of LAURDAN-stained live Vero cells under control conditions and following methyl-β-cyclodextrin (MβCD) treatment. **(A)** Zoomed spectral phasor plot showing the distributions of untreated cells (orange) and MβCD-treated cells (blue). The inset (top right) shows the full phasor plot circle, with the zoomed region delineated by a dashed rectangle. Reference locations corresponding to liquid order (Lo) and disorder (Ld) membrane components used for fraction analysis are indicated. **(B)** Histogram of the fractional contribution derived from component analysis, showing the distributions for the Lo fraction of untreated (orange) and MβCD-treated (blue) cells; dashed lines indicate the center of mass (CM) value for each condition. **(C–D)** Corresponding SFC measurements. **(C)** Zoomed spectral phasor plot of control untreated and MβCD-treated cells stained with LAURDAN, as well as untreated cells without LAURDAN to assess cell autofluorescence. The inset (top right) shows the full phasor plot circle, with the zoomed region indicated by a dashed rectangle. Reference locations corresponding to Lo and Ld membrane components, as well as the autofluorescence component, are indicated. **(D)** Histogram of contribution of the Lo fraction obtained from component analysis. Phasor distributions represent merged data from *n = 3* independent experiments per condition.

To define reference coordinates in phasor space for ordered and disordered membrane components, average phasor positions obtained from DOPC 0% cholesterol and DPPC 33% cholesterol MLVs were used (fluid to liquid-order linear combination), respectively, for each acquisition platform. These reference points enabled a two-component fraction analysis of LAURDAN emission in live cells.

Consistent with previous microscopy-based observations, HSI measurements revealed a clear green spectral displacement of LAURDAN fluorescence in the cellular phasor distribution following MβCD treatment relative to untreated controls. This result is indicative of an increase in membrane disorder associated with cholesterol depletion. Quantitatively, the center of mass of the liquid ordered fraction shifted from 0.36 in control cells to 0.30 following MβCD treatment, in agreement with earlier reports for the same cell type and experimental conditions (Díaz et al., 2026).

SFC measurements reproduced this separation in phasor space, with untreated and MβCD-treated cell populations occupying distinct regions. In this case, the center of mass of the liquid ordered fraction shifted from 0.40 in control cells to 0.35 following MβCD treatment. SFC measurements exhibited more compact distributions, with reduced dispersion compared to microscopy. One may notice that for SFC a phasor three-component analysis was used, this is due to the significant occurrence of autofluorescence picked up by SFC. In any case, phasor analysis performed as expected untangling the contribution of LAURDAN from liquid disorder or liquid order without contribution from autofluorescence.

### Inflammation-Associated membrane dynamic shift in bronchoalveolar macrophages measured by phSFC using LAURDAN fluorescence

To further challenge our phSFC approach we study the inflammation-associated membrane alterations *in vivo* using a mouse model of LPS-induced systemic inflammation (Burkard et al., 2023). Mice were treated with LPS or vehicle control, after which BAL was performed to isolate airway immune cells. BAL cells were stained with LAURDAN and antibodies against CD45 and CD11c, enabling selective analysis of macrophage populations by SFC (Figure 6).

**Figure 6.**
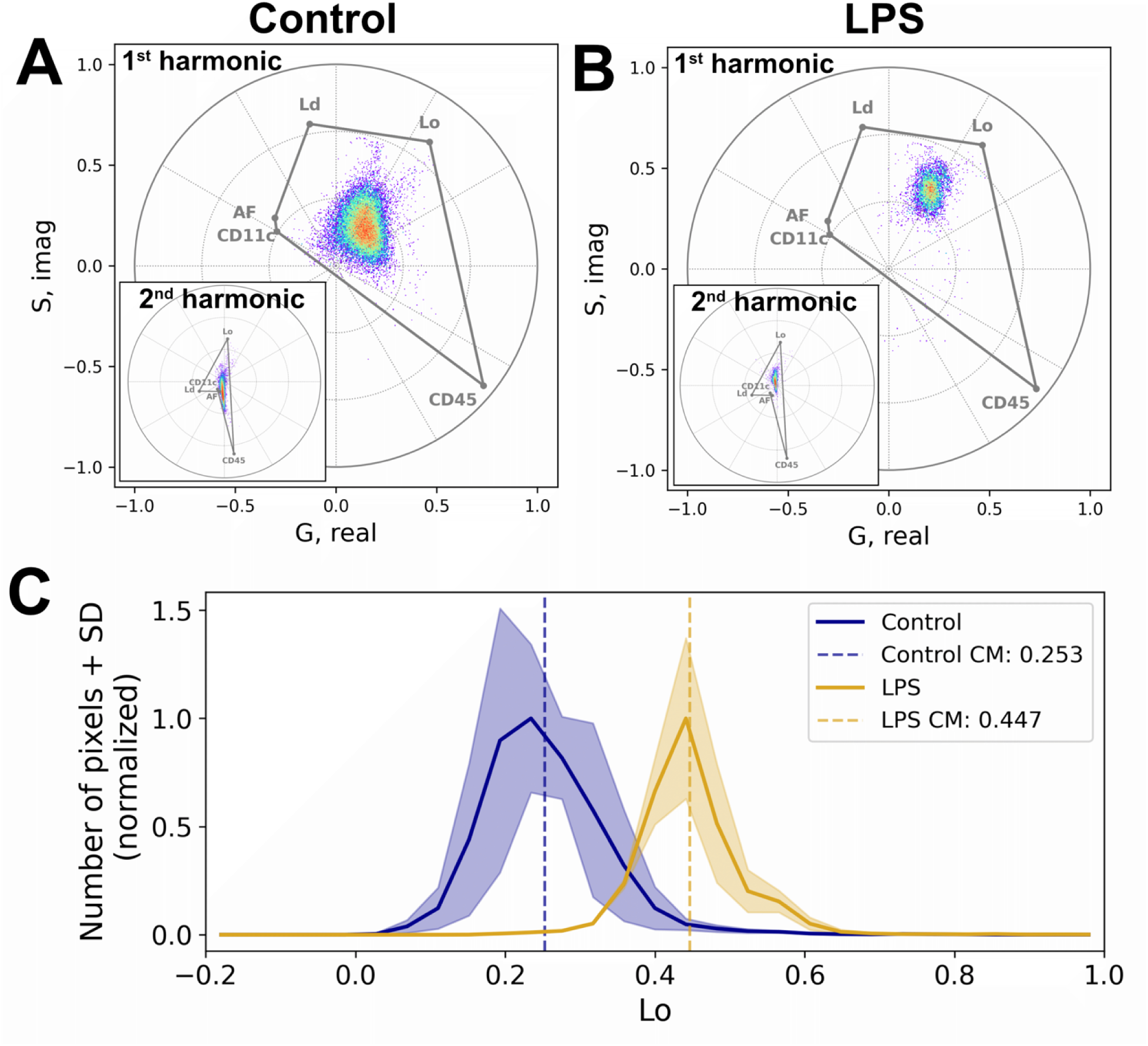
Phasor-based Spectral flow cytometry (phSFC) analysis of inflammation-induced membrane order changes in leukocytes from bronchoalveolar lavages. (**A, B**) Spectral phasor plots shown as two-dimensional histograms for bronchoalveolar lavage (BAL) leukocytes from **(A)** control mice and **(B)** LPS treated mice. The main panels display the first-harmonic spectral phasor plots, while the insets show the corresponding second-harmonic phasor plots. The phasor locations corresponding to the five components used for fractional analysis are indicated: LAURDAN fluorescence in liquid disorder (L_d_) or liquid order (L_o_) MLVs, CD45 (APC-Cy7 fluorescence), CD11c (eFluor-660 fluorescence) and autofluorescence (AF). (**C**) Mean distribution of the liquid order (L_o_) fraction by LAURDAN fluorescence was extracted from the five-component phasor analysis for control (blue) and LPS-treated (yellow) conditions. Shaded regions represent the standard deviation across biological replicates and the center of mass (CM) is indicated as a dashed vertical line. Data represent *n=3* for each condition.

Unlike the *in vitro* and cell culture experiments, analysis of BAL samples required consideration of multiple fluorescent contributors. In addition to LAURDAN emission from order and disorder membranes, CD45-APC-Cy7, CD11c-eFluor660 labeling and autofluorescence (AF) contributed to the detected spectral signal. To account for this complexity, a five-component phasor analysis was implemented. Phasor coordinates for autofluorescence, CD45-APC-Cy7 and CD11c-eFluore-660 were determined experimentally from unlabeled cells (for autofluorescence) or single-labeled beads, using both first and second harmonic components. This method (*n-harmonic* spectral phasor unmixing), previous develop by our group use reference coordinates for LAURDAN fluorescence from order and disorder membrane from DOPC 0% cholesterol and DPPC 33 % cholesterol MLVs, respectively, using the corresponding harmonic components (Vallmitjana et al., 2022).

Multi-component decomposition enabled extraction of the liquid order fractional at the single-cell level. Comparison of BAL leukocytes from control and LPS-treated mice revealed a clear liquid order increasement at the phasor distribution following LPS exposure (Figure 6E). Specifically, the fraction of the liquid-ordered LAURDAN fraction exhibited a displacement toward higher center-of-mass values in LPS-treated samples relative to controls (0.25 to 0.44 in control versus LPS-treated, respectively), indicating a more rigid membrane dynamics in cells due to inflammation-associated stimulus.

## Discussion

In this work, we establish an analytical framework for applying spectral phasor analysis to spectral flow cytometry (phasor-based Spectral Flow Cytometry, phSFC) that is directly comparable to the well-established HSI paradigm. By adapting the phasor approach to the distinct data structure and acquisition characteristics of flow cytometry, we enable fit-free spectral analysis at the single-cell level while preserving conceptual and interpretative continuity with microscopy-based phasor methods. This framework provides a common analytical basis for comparing spectral information across imaging- and flow-based modalities. Within this framework, we show that SFC can reliably reproduce LAURDAN-based membrane fluidity measurements traditionally obtained using HSI, while extending their applicability to high-throughput and complex biological settings. Across model membranes, live cells, and *in vivo* samples, phasor-based representations derived from both platforms were highly consistent, despite differences in optical configuration, spectral sampling, and detection strategies.

Using MLVs with defined lipid compositions, both HSI and SFC yielded the expected separation between fluid and order membrane phases in phasor space. While minor shifts in absolute cluster positions were observed between platforms, likely reflecting differences in spectral range and detector binning, the relative organization of lipid phases was preserved. Importantly, SFC measurements produced denser and more compact phasor distributions, consistent with the larger number of single-events sampled compared to pixel-based microscopy. An interesting aspect to discuss here is the signal-to-noise ratio when comparing phasor results from HSI and SFC. While in SFC the full MLV/cell contributes to the recorded spectrum, in HSI this information is spatially distributed due to the imaging resolution. This fact generates a main difference between SFC and HSI phasor representation. First, phasor clusters in SFC represent the behavior of a population of single events (MLVs or cells), where in HSI phasor clusters represent the heterogeneity of membrane dynamics due to spatial information from different regions of the MLVs or cells. Moreover, the signal-to-noise ratio differs due to the averaged fluorescence contribution in SFC, similar to bulk measurements in cuvette experiments using MLVs. In contrast, in HSI the contribution from different regions of the cell typically represents a small volume with a lower amount of fluorescence (signal from the point spread function, approximately 200–300 nm in x/y and 500–600 nm in z). Thus, information has to be compared carefully, but the two approaches are complementary in the sense of statistical representation (SFC) and spatio-temporal information (HSI).

Beyond quantitative comparison with microscopy, spectral phasor analysis in phSFC introduces an additional analytical dimension that can be directly exploited for event selection. Because the phasor representation encodes the full spectral signature of each event into a compact coordinate system, populations with distinct fluorescence properties naturally occupy well-defined regions of phasor space. This property enables phasor space to be used as an alternative gating domain, complementary to conventional scatter- or intensity-based gating strategies. As illustrated in Supplementary Figure S5, selecting events directly in phasor space allows populations with distinct spectral fingerprints to be readily identified and isolated without requiring prior knowledge of individual fluorescence channels.

This capability is particularly powerful in SFC, where complex fluorescence signatures arising from probes, autofluorescence, and antibody labeling often overlap in conventional parameter space. In contrast, the phasor representation provides an intuitive and physically meaningful organization of spectral information, in which related spectral signatures cluster together and mixtures fall along predictable trajectories. As a result, phasor-based gating can reveal spectroscopic defined subpopulations that may be difficult to identify using standard gating strategies alone.

The use of phasor space as an additional gating dimension therefore extends the utility of phSFC beyond quantitative membrane analysis, providing a general framework for spectrally informed population selection. This capability is particularly relevant for spectral flow cytometry applications involving biosensors, such as FRET- or calcium-based probes, or other sensors that exhibit activity-dependent spectral shifts (Foncillas et al., 2024; Henderson et al., 2022; Piatkevich and Verkhusha, 2011). When combined with conventional gating approaches, phasor-based selection offers a flexible strategy for exploring complex datasets and identifying biologically meaningful subpopulations based on their intrinsic fluorescence signatures.

Analysis across increasing cholesterol concentrations further supported the uniqueness of phasor analysis for LAURDAN fluorescence in membranes and the agreement between platforms. For DOPC membranes, both modalities captured a progressive, approximately linear shift in phasor space consistent with a linear combination of spectral components. The phasor blue shift is consistent with the expected increase in membrane order due to the cholesterol percentage. Despite the blue shift observed in the phasor plot for DOPC membranes (smaller phase values), they also exhibit lower modulation, consistent with a broader LAURDAN spectrum arising from the more heterogeneous environment experienced by the probe in MLVs.

In contrast, DPPC and mixed DOPC/DPPC membranes exhibited non-linear phasor trajectories with cholesterol addition, reflecting the more complex phase behavior of saturated and mixed lipid systems. We have demonstrated this complex behavior in previous studies using cuvette experiments and MLVs with similar compositions (Malacrida, 2014). These deviations were consistently detected by both HSI and SFC, underscoring the sensitivity of spectral phasor analysis to non-ideal membrane organization. The results obtained with MLVs reinforce our previous findings demonstrating the advantages of spectral phasor analysis over the generalized polarization (GP) function for LAURDAN and other solvatochromic membrane probes (Malacrida et al., 2015). In contrast, GP analysis of spectral data would yield intermediate values between the two components, effectively averaging out the rich photophysics of LAURDAN imposed by the GP mathematical model (Andronico et al., 2025).

Re-representation of the data to explicitly test linear combination predictions revealed that phasor distributions obtained by SFC were more closely centered around theoretical intermediate values and displayed reduced dispersion compared to microscopy-derived distributions. This increased convergence toward expected phasor positions reflects the strong statistical averaging afforded by flow-based acquisition as discussed previously.

Extending the approach to live cells, both HSI and SFC detected cholesterol-dependent shifts in LAURDAN phasor distributions following methyl-β-cyclodextrin treatment, consistent with changes in membrane order. MβCD has been used extensively to modify membrane dynamics (Mahammad and Parmryd, 2015). Empty MβCD can be used to remove cholesterol from membranes, affecting membrane order in different regions of the cell. Cholesterol removal by MβCD generates a gradient imbalance between the plasma membrane and intracellular membranes that depends on the incubation time (Rouquette-Jazdanian et al., 2006). Here, we used a standard endpoint previously established by our group that induces a strong imbalance in cholesterol between plasma membrane and intracellular membrane fluidity/order (Díaz et al., 2026). phSFC was able to capture the average change in membrane order fraction in high-statistical single-cell experiment (around 20000 cells per experiment) in consistently with HSI results. This comes at the cost of losing spatial and temporal information.

While the overall direction of the phasor shifts was conserved across modalities, cytometry-derived measurements exhibited more compact distributions. This highlights the advantage of single-cell-resolved, high-throughput sampling for resolving heterogeneous responses and detecting treatment-induced shifts that may be spatially diluted or unevenly represented in microscopy measurements. Finally, our results in BAL macrophages are consistent with previous reports showing that changes in plasma membrane organization and fluidity accompany inflammatory activation (Adams et al., 2014; de la Haba et al., 2016; Printen et al., 1993). However, it is noteworthy that our observations of an increased liquid-order fraction (higher GP) appear to diverge from certain *in vitro* studies, such as those by (de la Haba et al., 2016). Using LAURDAN-based microscopy on THP-1 cell lines, that study reported that LPS stimulation induces a “fluidification” (lower GP) of the plasma membrane to facilitate TLR4-dependent signaling and TNF-α secretion. Our contrasting findings, a shift toward a more order state in primary BAL macrophages, may be explained by several factors. First, primary alveolar macrophages exist in a unique lipid-rich pulmonary environment and possess a distinct lipidome compared to immortalized monocytic lines like THP-1, which may alter their structural response to LPS. Second, while microscopy often focuses exclusively on the plasma membrane, our phSFC approach captures the “whole-cell” LAURDAN signal. This likely includes the expansion and rigidification of internal secretory organelles, such as the Endoplasmic Reticulum and Golgi, which undergo massive structural reorganization to support cytokine production during activation. Lastly, the timing of the measurement may reflect different phases of activation; an initial fluidification for receptor recruitment may be followed by a sustained increase in order as signaling micro-domains coalesce. Moreover, the previously discussed differences between spectral phasor and GP analysis of LAURDAN fluorescence may partly explain the discrepancy between our results and those reported by de la Haba *et al*. 2016.Importantly, our data extend these observations by demonstrating that LAURDAN-based membrane readouts can be robustly detected at the single-cell level using phSFC, providing a complementary and scalable alternative to microscopy-based approaches for studying inflammation-associated membrane dynamics in heterogeneous cell populations.

An important aspect to highlight about our approach (phSFC) is that it can handle multilabeling, including autofluorescence, by analyzing the data using phasor-based multicomponent analysis (Vallmitjana et al., 2022). This approach is particularly key for phSFC, where multiple labeling is commonly used to molecularly fingerprint the analyzed cell populations. In our case, we were able to handle the complex spectroscopy of LAURDAN fluorescence simultaneously with autofluorescence and an antibody-derived signal to define the cellular group of interest in BAL. Depending on the needs of the study, one can extend this multilabeling approach to up to five different components plus the LAURDAN signal using higher harmonic information (second harmonic for three components plus LAURDAN and third harmonic for five components plus LAURDAN) (Vallmitjana et al., 2022). Nevertheless, careful monitoring of the signals is required to avoid poor signal-to-noise ratios, where higher harmonic phasor analysis will perform poorly.

Together, these results establish SFC as a robust and scalable extension for LAURDAN phasor analysis, maintaining direct continuity with established HSI approaches while enabling quantitative membrane biophysical measurements in complex cellular and *in vivo* systems. At the same time, our results emphasize the complementary nature of SFC and microscopy-based spectral phasor analysis. While SFC provides exceptional statistical robustness, population-level resolution, and sensitivity to subtle spectral shifts across large numbers of cells, it inherently lacks spatial context within individual cells and tissues, including subcellular heterogeneity and spatial organization of membrane properties. In contrast, HSI preserves this spatial information, enabling direct visualization of membrane order variations within and across cellular compartments, albeit at the cost of lower throughput and increased sensitivity to imaging-related artifacts.

Viewed together, these modalities form a synergistic analytical toolkit. HSI can be leveraged to uncover spatially resolved biophysical mechanisms and guide hypothesis generation, while SFC enables rapid, unbiased validation and quantification of these findings across several orders of magnitude of single-cell events in heterogeneous cell populations. By establishing a unified phasor-based analytical framework applicable to both platforms, this work facilitates direct cross-modal comparisons and supports integrated experimental strategies that combine spatial insight with statistical power. Such combined approaches are particularly well suited for interrogating complex biological processes, including membrane remodeling during immune activation, metabolic transitions, and disease-associated cellular heterogeneity.

## Acknowledgments

The authors also acknowledge using OpenAI’s ChatGPT (version GPT-4, accessed February 2026) exclusively for language refinement and style correction; all AI-assisted edits were reviewed and approved by the authors.

## Ethical Approval

All animal procedures were conducted in accordance with national and institutional guidelines for the care and use of laboratory animals. The experimental protocol was reviewed and approved by the National Ethics Committee for Animal Use under the animal use protocol number 2425.

## Author Contributions

Conceptualization: LM, BP, PC

Methodology: BP, PC, MD, DA, AR, LM

Software: BP

Formal Analysis: BP, PC

Investigation: BP, PC, MD, DA, AR

Resources: LM

Data Curation: BP,PC, AR, LM

Writing – Original Draft: BP, PC

Writing – Review & Editing: MD, LM

Visualization: BP

Supervision: LM

Project Administration: LM

Funding Acquisition: ML

All authors have read and approved the final version of the manuscript.

