## Supplementary Material for "High-Throughput Single-Cell Spectroscopy Using Phasor Analysis of Spectral Flow Cytometry"

­

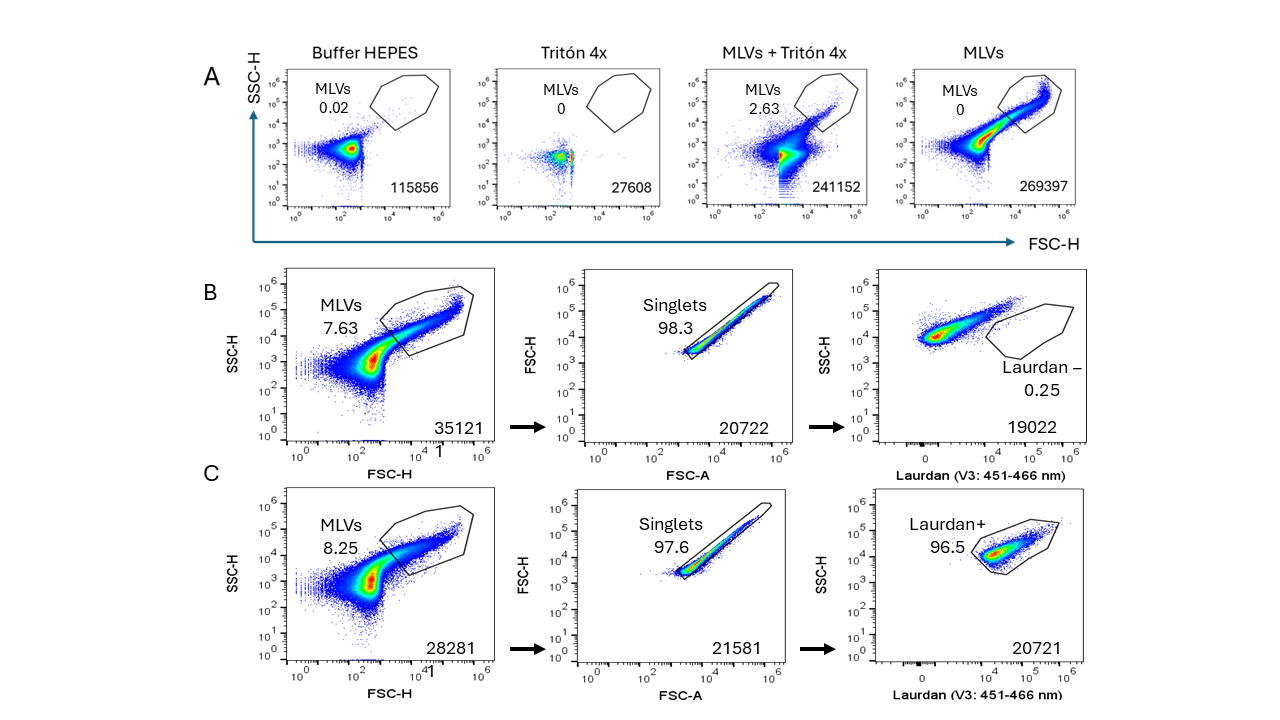

LAURDAN +

LAURDAN -

**­­­­**

**Supplementary Figure S1. Gating strategies for SFC analysis of MLVs.** **(A)** Control samples used to validate the correct identification of MLVs population and to exclude electronic noise and debris. HEPES buffer alone was acquired to define the electronic noise threshold. A 4X Triton solution was acquired to assess the effect of detergent-mediated lipid solubilization. In addition, MLVs treated with 4X Triton were acquired to confirm that the gated population corresponded to lipid vesicles rather than noise or aggregates. The MLVs sample used for these controls consisted of a DPPC:DOPC (1:1) mixture with 33% cholesterol and is shown in the last dot plot of this row. **(B)** Representative gating strategy for MLVs, including SSC-A versus FSC-A for initial population definition and FSC-A versus FSC-H for singlet selection. **(C)** Identification of LAURDAN-negative and LAURDAN-positive MLVs populations based on fluorescence intensity in V3 channel. For each gating step, the number of events within the gate and the corresponding frequency of each population are displayed. LAURDAN fluorescence was analyzed across all 16 channels of the violet laser. For simplicity and clarity, representative plots display fluorescence intensity in the V3 channel, which corresponds to the maximum emission of LAURDAN.

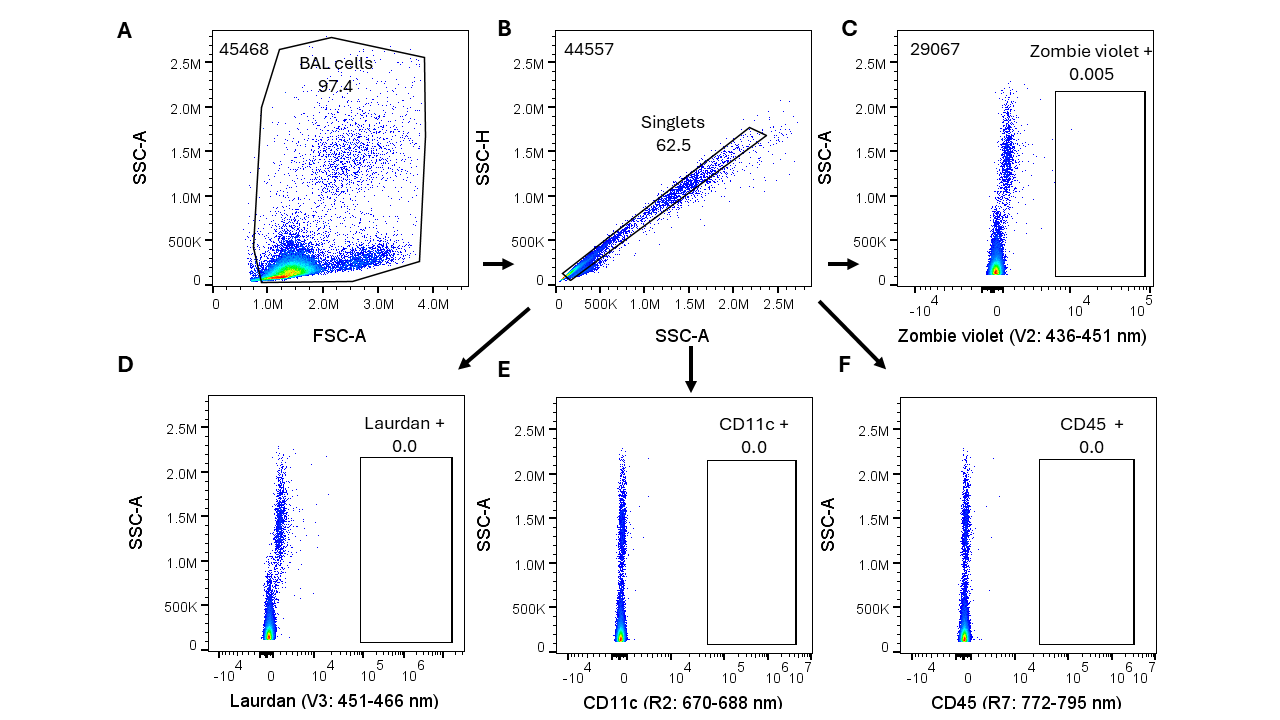

LAURDAN +

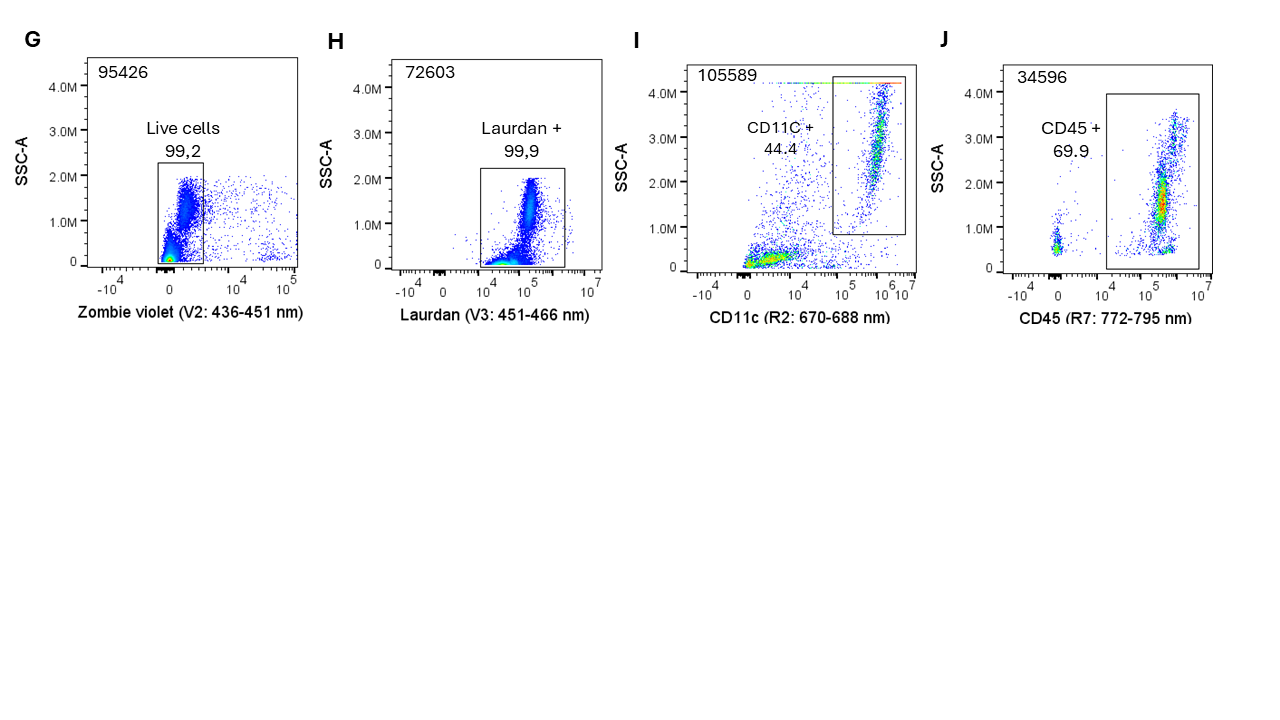

LAURDAN +

**Supplementary Figure S2. Controls and gating strategy for antibody panel selection in BAL samples. (A)** Forward scatter area (FSC-A) versus side scatter area (SSC-A) used to identify and gate the leukocyte population. **(B)** FSC-A versus FSC-H gating was performed to exclude doublets and select singlet events. Based on the singlet population, negative control samples for each fluorescent marker included in the staining panel are shown: **(C)** Zombie Violet viability dye, **(D)** LAURDAN, **(E)** CD45 APC-Cy7, and **(F)** CD11c eFluor 660. These controls were used to define negative populations and establish accurate gating boundaries for subsequent immunophenotyping analyses. For each gating step, the number of events within the gate and the corresponding frequency of each population are displayed. LAURDAN fluorescence was analyzed across all 16 channels of the violet laser. For simplicity and clarity, representative plots display fluorescence intensity in the V3 channel, which corresponds to the maximum emission of LAURDAN.

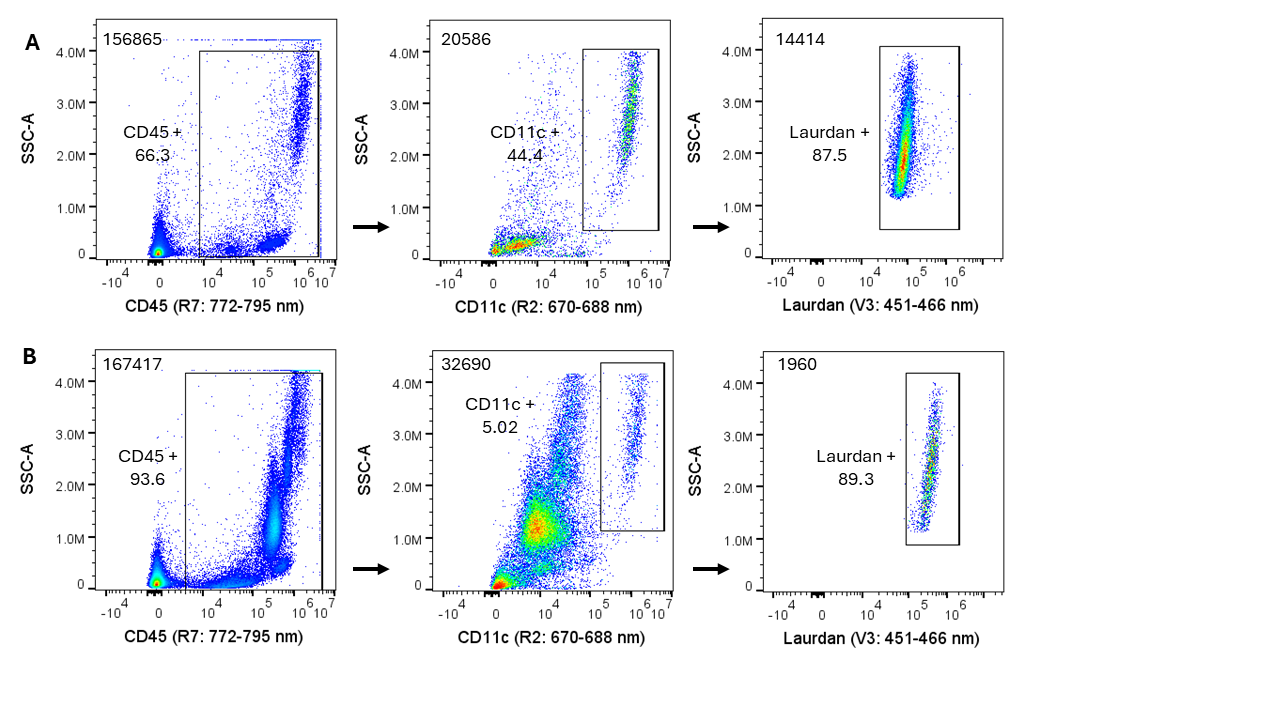

LAURDAN +

LAURDAN +

**Supplementary Figure S3. Gating strategy for identification of LAURDAN-positive macrophages in BAL samples. (A)** Representative gating strategy applied to bronchoalveolar lavage (BAL) samples from a control animal to identify macrophages defined as CD45⁺ CD11c⁺ cells that are positive for LAURDAN. **(B)** The same gating strategy applied to BAL samples from an LPS-treated mouse. For each gating step, the number of events within the gate and the corresponding frequency of each population are displayed. LAURDAN fluorescence was analyzed across all 16 channels of the violet laser. For simplicity and clarity, representative plots display fluorescence intensity in the V3 channel, which corresponds to the maximum emission of LAURDAN.

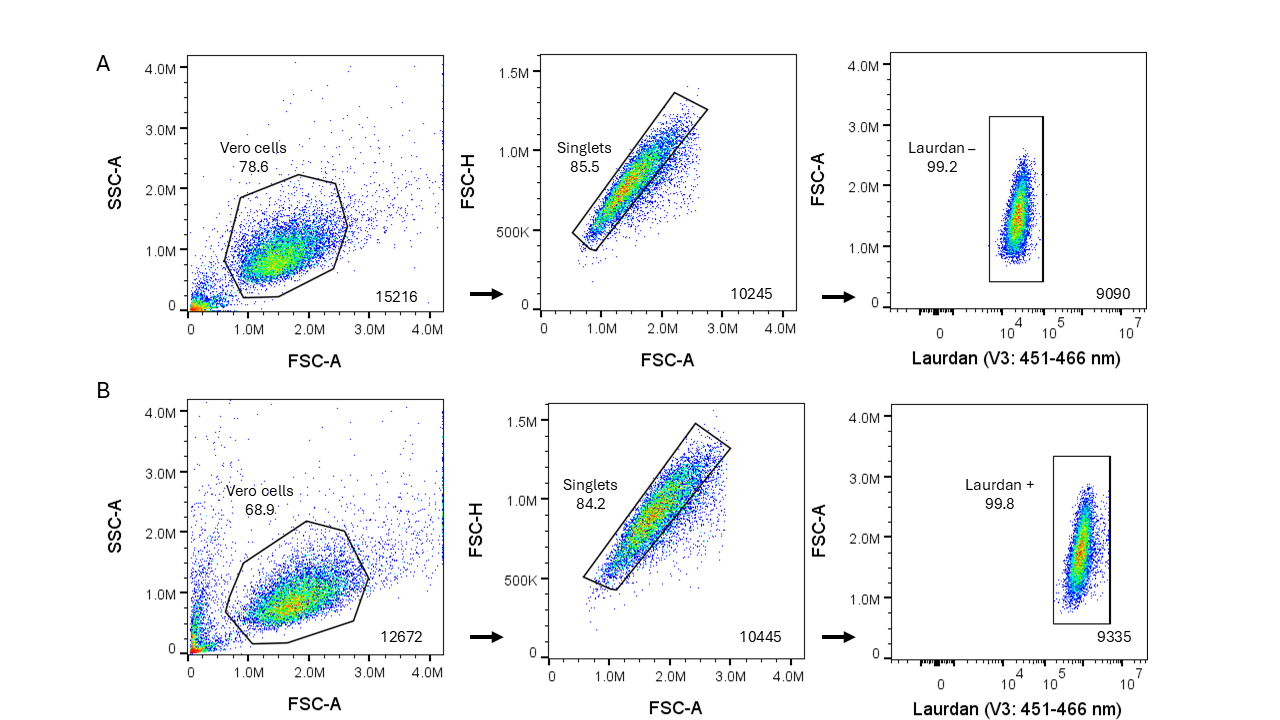

**LAURDAN**

**LAURDAN**

LAURDAN +

LAURDAN -

**Supplementary Figure S4. Gating strategy for analysis of Vero cells. (A)** Gating strategy used to identify Vero cells, including selection of the Vero cell population based on SSC-A versus FSC-A, exclusion of doublets using FSC-H versus FSC-A, and identification of LAURDAN-negative cells. **(B)** The same gating strategy applied to identify LAURDAN-positive VERO cells. For each gating step, the number of events within the gate and the corresponding frequency of each population are displayed. LAURDAN fluorescence was analyzed across all 16 channels of the violet laser. For simplicity and clarity, representative plots display fluorescence intensity in the V3 channel, which corresponds to the maximum emission of LAURDAN.

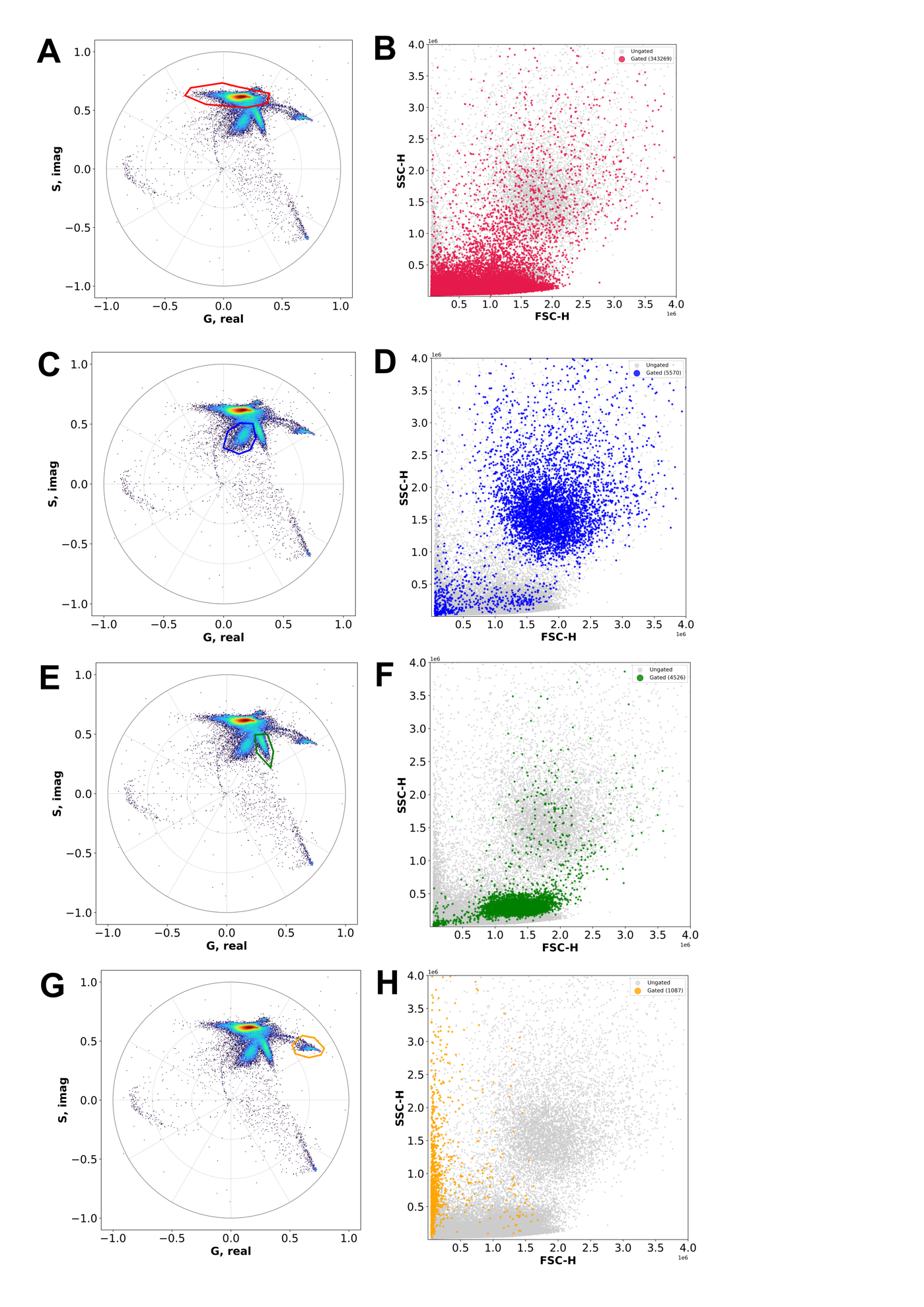

**Supplementary Figure S5. Enhanced population discrimination through phasor space gating of an ungated control BAL sample.** Spectral phasor plots and corresponding scatter plots illustrating the use of phasor space as an alternative gating domain in phSFC. All panels show data from the same control BAL sample with no prior gating applied, so that all acquired events are included. Panels A, C, E, and G show spectral phasor distributions with four different regions selected in phasor space, each indicated by a distinct colored gate. Panels B, D, F, and H show the corresponding SSC-H vs FSC-H dot plots for the same dataset, where all events are displayed in gray and events falling within the respective phasor gate are highlighted in the matching color. This representation illustrates how phasor space provides an additional domain for event selection based on spectral fluorescence signatures, allowing populations with distinct emission characteristics to be identified independently of conventional scatter-based gating. The correspondence between phasor-selected events and their distribution in scatter space demonstrates the potential of phasor-based gating to reveal spectrally defined subpopulations in complex samples.
